# Experimental evolution of *Candida albicans* under hypoxia and heat shock reveals nuclear genome variants and mitochondrial methylome alterations

**DOI:** 10.1101/167338

**Authors:** Thais Fernanda Bartelli, Danielle do Carmo Ferreira Bruno, Flavio Lichtenstein, Marcelo R. S. Briones

## Abstract

Infection by *Candida albicans* requires its adaption to physical constraints in the human body, such as low oxygen tension (hypoxia), increased temperature (37°C) and different carbon sources. Previous studies demonstrated that the genetic variability of *C. albicans* isolates is an important adaptive mechanism, although little is known about the dynamics of this genetic diversity, and the influence of these environmental conditions on its mitochondrial genome (mtDNA). To test the synergistic effect of these stress conditions on *C. albicans* genome, reference strain SC5314 was subjected to an *in vitro* evolution scheme under hypoxia and 37°C, with two different carbon sources (glycerol and dextrose) for up to 48 weeks (approximately 4,000 generations). Experimental evolution results showed no sequence or copy number changes in the mtDNA, although sequence variants were detected on its nuclear genome by Multilocus sequence typing (MLST) and whole genome sequencing (WGS). After 12 weeks of experimental evolution, sample GTH12, grown under hypoxia at 37°C in glycerol, showed inferior growth and respiratory rates as compared to other conditions tested. Although WGS of GTH12 revealed no variants in its mtDNA, WGS with sodium bisulfite showed a significant reduction in mtDNA methylation in GTH12 in both non-coding and coding regions. Our results provide the first whole mitochondrial genome methylation map of *C. albicans* and show that environmental conditions promote the selective growth of specific variants and affect the methylation patterns of the mtDNA in a strain-specific manner.

## INTRODUCTION

Host-microbe interactions are dynamic processes, where microbes are constantly shaping their genotypic and phenotypic responses according to the host’s physiological conditions for successful colonization and infection. Compared to bacteria, fungi are a small part of the human microbiome (0.1%) (Qin et al., 2010; Underhill and Iliev, 2014; Arumugam et al. 2011), *C. albicans* is ubiquitously found in the human body, and can successfully colonize diverse niches, such as skin, urogenital and gastrointestinal tracts, including internal organs after tissue invasion and bloodstream dissemination (reviewed in Brock, 2009; Huffnagle and Noverr, 2013; Underhill and Iliev, 2014). Although a part of the human microbiota, *C. albicans* causes severe mucosal and bloodstream opportunistic infections in immunosuppressed hosts, with nearly 400,000 nosocomial cases worldwide with 46 to 75% mortality rates (Brown et al., 2012). *C. albicans* commensal or pathogenic outcomes can be influenced by interactions of the fungus with other components of the resident microbiota, such as gram-positive bacteria, including synergism or antagonism (Förster et al., 2016). At least 1 in 4 patients (25%) with positive *C. albicans* bloodstream infection also have one or more bacterial species associated (Klotz et al., 2007) and polymicrobial bloodstream infections are commonly associated with coagulase-negative Staphylococci, usually *S. epidermidis*, and *Candida* species (Karlowicz et al., 2002; Pammi et al., 2013; Raad and Hanna, 2002; Sutter et al., 2008).

The metabolic plasticity of *C. albicans* allows its survival and response to multiple environmental stimuli simultaneously, such as temperature, oxygen and nutrient availability. These environmental factors can diverge depending on the complexity of the human niche colonized (Brown et al., 2014) and is different from those typically used by researchers or clinicians in the laboratory environment, which tends to favor *C. albicans* optimal growth rather than replicate the host physiological conditions. Whilst the oxygen tension in the atmosphere is close to 21% (or an O_2_ pressure of 159 mmHg at sea level), known as normoxia, healthy tissues in the human body are expected to have O_2_ levels of 2.5 to 9% (Grahl et al., 2012). This lower oxygen tension (< 21%), known as hypoxia, varies depending on the anatomical site or the presence of tissue inflammation (Grahl et al., 2012). *C. albicans* is also able to adjust to temperature fluctuations. Above 30°C the heat shock response is activated, leading to transcription of several genes, especially chaperones, by the heat shock factor 1 (Hsf1) (Nicholls et al., 2009). In the non-pathogenic yeast *Saccharomyces cerevisiae*, the heat shock transcription factor (Hsf) changes its conformation in response to superoxide and temperature (Lee et al., 2000), while the transcription factor Efg1, in *C. albicans*, acts in the hypoxia response, regulating the transcription of metabolic genes and also in the yeast-to-hyphal transition in a temperature dependent manner. When under hypoxia, this transcription factor inhibits the filamentous growth at temperatures ≤ 35°C, whilst at 37°C it is essential for this transition, independent of normoxia or hypoxia environments (Desai et al., 2015; Setiadi et al., 2006). While the yeast cell responses to heat shock and hypoxia can be related and dependent, many studies also show that changes in the carbon source strongly influence *C. albicans* stress resistance, and it is important to consider the pathogen response to combinatorial stressors rather than to individual stress factors commonly studied separately *in vitro* (Brown et al., 2014). The combinatorial stresses are relevant when representing many host niches, such as a mucosal invasion (oxidative stresses plus water balance) or a kidney infection (cell adaptation to high salt concentrations plus endogenous reactive oxygen species), for instance (Brown et al., 2014).

Several studies have shown high intraspecific genetic variability of *C. albicans*, suggesting a correlation with adaptation and microevolution when colonizing the human body (Forche et al., 2009; Morschhäuser et al., 2000; Wartenberg et al., 2014). This genetic diversity is characterized by point mutations, amplification or deletion of chromosomal fragments, translocations, inversions and aneuploidy (Forche et al., 2011; Odds et al., 2006; Wartenberg et al., 2014). Besides genetic diversity, few studies indicate the distribution and significance of *C. albicans* epigenetic plasticity as an adaptive mechanism (Freire-Benéitez et al., 2016; Lopes da Rosa and Kaufman, 2012; Tscherner et al., 2015; Zordan et al., 2006). To date, the few studies on *C. albicans* epigenetic regulation focus on the epigenetics of its nuclear genome, mainly on the modulation of its morphology and other virulence factors, such as white-opaque switching (Freire-Benéitez et al., 2016; Kim et al., 2015; Mishra et al., 2011; Tscherner et al., 2015; Zhang et al., 2013; Zordan et al., 2006). There is increasing evidence of mtDNA methylation role in several diseases and its potential use as a biomarker for harmful environmental and nutritional factors (Iacobazzi et al., 2013). Mapping of *C. albicans* nuclear genome hypermethylated sites identified genes involved in morphogenesis and hyphal growth (16.7%), white-opaque switching (3.3%), iron use (6.7%), drugs resistance and signaling (12%), stress response (7.3%), and genes involved in regulatory activities such as chromatin organization (3.3%), cycle or cell division (7.3%), biogenesis and protein transport (12.7%), DNA / RNA processing (5.3%), pathogenesis or virulence (2%), carbohydrate metabolism (1.3%), and genes with unknown functions (22%). It was also observed that, in this species, methylation occurs at both CpG and CH sites (H = Adenine, Cytosine or Thymine), and mainly in the gene bodies, instead of the promoters. Mapping of hypermethylated sites in *C. albicans* genome revealed that the majority were within genes (82%), followed by non-repetitive intergenic regions (13%) and repetitive regions (5%), suggesting that methylation in this yeast genome plays an important role in gene regulation (Mishra et al., 2011). However, to date, there are no studies on *C. albicans* mitochondrial genome methylation.

Although an increasing amount of data points mitochondria as a key player in yeast virulence and survival (Bambach et al., 2009; Qu et al., 2012; Thomas et al., 2013), still little is known about *C. albicans* mtDNA variability and how host environmental factors influence its dynamics. A long-term ongoing experiment with *Escherichia coli*, by Richard Lenski group since 1988, has experimentally evolved 12 initially identical *E. coli* populations through over 50,000 generations and has given major insight into genetic aspects involved with bacterial evolution (Lenski, 2011). Although experimental evolution protocols are of great benefit and provide insights into evolutionary processes, only a few authors have considered this method for *C. albicans* so far. There are limited studies focused on the evolution of drug resistance (Cowen et al., 2000; Huang et al., 2011) or models with pathogen-host co-evolution (Cheng et al., 2007; Forche et al., 2009; Wartenberg et al., 2014). Understanding *C. albicans* tempo and mode in evolution using *in vitro* models are promising not only due to its medical relevance, but because *C. albicans* has the potential to emerge as a model organism for fungal-pathogen interaction studies, along with *Aspergillus fumigatus* and *Cryptococcus neoformans* (Kabir et al., 2012). These data could help bring light into *C. albicans* adaptations to its host and understanding of its virulence.

In the present study, *C. albicans* reference strain SC5314 was continuously exposed to hypoxia and 37°C in a natural polymicrobial experimental evolution scheme aiming to investigate the synergistic effect of these stress factors on *C. albicans* mtDNA. Our data showed differential growth in different temperatures, oxygen and nutrient combinations and the occurrence of nuclear but not mitochondrial sequence microvariations. Although we have not identified sequence or copy number changes in the mtDNA, we mapped, for the first time, the methylation sites in *C. albicans* mitochondrial genome. Another strain, L757, a more recent pathogenic isolate (Padovan et al., 2009), was analysed in parallel. Our results indicate that environmental conditions, such as continuous exposure to hypoxia and 37°C in a polymicrobial culture, which simulates the human body, affect the methylation patterns of this genome in a strain-specific manner.

## RESULTS

### Experimental evolution of strain SC5314

For experimental evolution, *C. albicans* strain SC5314 was continuously maintained in normoxia at 28°C or hypoxia at 37°C, in rich dextrose (YPD) or non-fermentable glycerol media (YPG) for a period of 48 weeks (or 3,000 to 4,000 generations depending on growth conditions). Samples were named accordingly (Table 1) and different time points were analyzed, namely P0 (yeasts in pre-experimental evolution, grown for 48h) or P1, P12, P24 and P48 which were continuously grown for 1, 12, 24 or 48 weeks, respectively.

**Table 1.** Growth conditions used in the experimental evolution of *C. albicans* strain SC5314. Samples were named according to the oxygen (normoxia = N or hypoxia = H), temperature (37°C = T) and carbon source (YPD medium = D, YPG medium = G) followed by the number of weeks they were grown (1, 12, 24 or 48 weeks).

| Oxygen level | Temperature | Media | Nomenclature |
| --- | --- | --- | --- |
| Normoxia | 28 °C | YPD | DN |
|  |  | YPG | GN |
| Hypoxia | 37 °C | YPD | DTH |
|  |  | YPG | GTH |

Coisolation of *C. albicans* and *Staphylococcus epidermidis* is frequent and tight commensal interactions between these two species are observed in the human body, the experimental evolution scheme was carried out with polymicrobial cultures to investigate *C. albicans* genome evolution in conditions that approximated a host-simulating environment. Surprisingly, sequencing of sample GTH 16S rRNA after 48 weeks of experimental evolution revealed the existence of a second bacterial species in our yeast cultures. While in yeast cultures of GTH1 and GTH12 we could only identify *S. epidermidis* by 16S rRNA and *tuf* sequencing, we identified the slow growing anaerobic gram-positive bacteria *Propionibacterium acnes* in the GTH48 but not in the DVN (P1, P12, P48) cultures.

Yeast doubling times (Dt) were estimated for samples P1, and data revealed increased Dt for GN and GTH, indicating smaller growth rate for *C. albicans* SC5314 when glycerol is the carbon source available. Samples exhibited different Dt, with significant differences between GN1 (Mean ± SEM 3.53 ± 0.15) *versus* DN1 (Mean ± SEM 2.38 ± 0.12) and GTH1 (Mean ± SEM 4.58 ± 0.41) *versus* DTH1 (Mean ± SEM 2.97 ± 0.14) comparisons (Figure 1A). Sample GTH1 also showed higher Dt when compared to DN1, which is the common environmental lab condition usually applied to *C. albicans* cultures, i.e. normoxia, room temperature and dextrose-rich medium.

**Figure 1.**
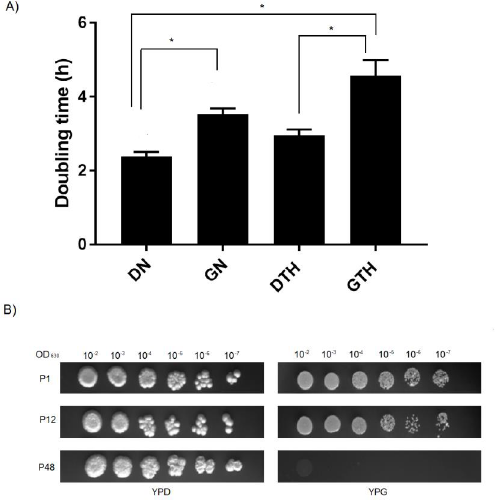
*C. albicans* SC5314 growth under different oxygen, temperature and nutrient combinations. **A)** Doubling times for *C. albicans* SC5314 grown for one week (P1). Plates were incubated at normoxia/28°C (DN, GN) or hypoxia/37°C (DTH, GTH) and the OD_630nm_ was measured at 5h, 10h and 24h intervals. Data shown as mean ± SEM of three independent experiments in triplicate. Statistically significant differences (P < 0.05) are denoted by letters (a, b and c). **B)** Spot assay for samples DN and GTH, showing a lack of yeast cells after 48 weeks of experimental evolution for sample GTH.

Yeast cell viability was checked by counting of viable cells (P1, P12 and P48) in the presence of Trypan Blue in optical microscopy. Cell viabilities for all samples were over 86% and cells grew adequately during the 48 weeks period, with the exception of sample GTH. As confirmed by cell counting on hemocytometer chamber and an attempt of *C. albicans ACT1* amplification from culture DNAs, yeasts were extinct from the culture around 24 weeks after experimental evolution initiation (data not shown), therefore precluding any analysis after this point (i.e. GTH48) (Figure 1B).

Samples achieved and average of 52 (P1), 630 (P12) and 2777 (P48) generations, as calculations inferred by the Dt. In normoxia and dextrose, *C. albicans* showed the highest number of generations, i.e. DN (P1= 70, P12= 840, P48= 3360), followed by DTH (P1= 56, P12= 672 and P48= 2688), GN (P1= 47, P12= 571, P48= 2284) and GTH (P1= 36, P12=436, P48= n/a).

In low oxygen conditions, microorganisms need to adjust their metabolism, increasing respiration or using alternative mechanisms, such as fermentation. We estimated the respiratory rate of samples by measuring the oxygen consumption in three different time points (P0, P12, P48) (Figure 2). In 48h hypoxia or normoxia (P0) samples showed similar respiratory rates, with the exception of DN (Mean ± SEM 3.34 ± 0.16), which showed significant smaller value as compared to DTH (Mean ± SEM 4.96 ± 0.28). Comparison of oxygen consumption through time shows that values were maintained for those cultivated under normoxia, that is DN (Mean ± SEM P0= 3.34 ± 0.16, P12= 2.80 ± 0.38, P48= 2.90 ± 0.21) and GN (Mean ± SEM P0= 3.97 ± 0.15, P12= 3.83 ± 0.16, P48= 3.61 ± 0.30). On the other hand, there was a reduction in the respiratory rate for samples cultivated in hypoxia, i.e. DTH (Mean ± SEM P0= 4.96 ± 0.28, P12= 4.40 ± 0.45, P48= 3.38 ± 0.34) and GTH (Mean ± SEM P0= 4.53 ± 0.45, P12= 1.85 ± 0.45, P48= n/a), with a significant reduction between DTH48 *versus* DTH and GTH12 *versus* GTH. In addition to time, the carbon source also seems to be an important determinant of the respiratory rate when in hypoxia. By comparing samples DTH and GTH, we can see that the yeast tends to consume more oxygen when in the presence of dextrose, with a significant respiratory rate difference between samples DTH12 and GTH12 (Figure 2).

**Figure 2.**
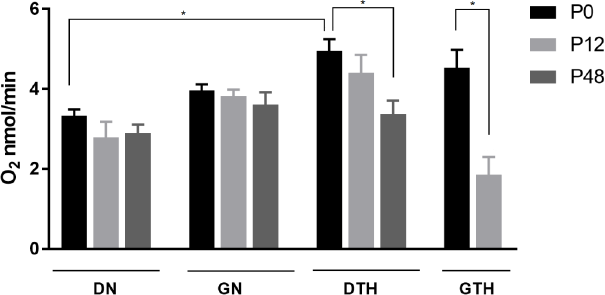
Oxygen consumption of *C. albicans* SC5314 grown for 48h (P0), 12 (P12) or 48 (P48) weeks under different oxygen and temperature combinations. Data shown as mean ± SEM of O_2_ (nmol/min) intake by 2×10^6^ cells in three independent experiments in triplicate. (*) statistically significant differences (P < 0.05). It was not possible to perform O_2_ measures for sample GTH48 due to an insufficient amount of yeast cells on culture. Samples under normoxia/28°C (DN, GN) and hypoxia/37°C (DTH, GTH).

### *In vitro* exposure to hypoxia and heat shock does not change C. *albicans* mtDNA sequence and copy number

To investigate if the long-term exposure to hypoxia and 37°C (physical conditions in the human body) could influence the intraspecific sequence diversity among the mtDNA of *C. albicans*, we sequenced three mtDNA non-coding regions and a complex III chain gene, cytochrome B (COB), for the SC5314 strain throughout the *in vitro* evolution. Sequencing of samples DN, GN, DTH and GTH (Table 1) mtDNA intergenic regions, namely IG1 (tRNA-Gly/*COX1*), IG2 (*NAD3*/*COB*) and IG3 (RRNS/*NAD4L*) (Bartelli et al., 2013), showed no sequence variability in experimental samples mtDNA as compared to sample P0 after 6, 12, 24 or 48 weeks of growth. These same samples showed no nucleotide changes for *COB* after 24 or 48 weeks of experimental evolution. The successful amplification of long amplicons, such as Long 2 (10.2kb amplicon from 13105 to 23210bp, between *ATP6* and *NAD3*) and Long 3 (10.3kb amplicon from 23210 to 33578 located between *NAD3* e *NAD4*) confirmed the non-existence of mtDNA sequence variability in these samples (data not shown). The absence of smaller unspecific bands and efficient amplification of the long amplicons after 12 and 48 weeks, confirms the lack of significant sequence variation and/or lesions in the mtDNA that could affect the DNA polymerase progression throughout the DNA molecule (data not shown).

To test if the different oxygen and temperature combinations could influence the amount of mtDNA per cell, we have quantified the number of mtDNA molecules for the samples P0, P12 and P48. The mean number of mtDNA molecules per diploid nuclear genome was measured by *COX2* Real time PCR quantification normalized by the nuclear *ACT1* (Figure 3). Samples grown for 48h under normoxia at 28°C or hypoxia at 37°C showed no difference between mtDNA copy number per cell among them (Table 2), and the amount of mtDNA in the samples remained stable throughout the experimental evolution (Figure 3). In week 12 of experimental evolution, samples DN12 (Mean ± SEM: 52.7 ± 9.8) showed increased amount of mtDNA copies/cell as compared to samples cultured for the same period with glycerol (GN12) (Mean ± SEM: 16.7 ± 2) or in hypoxia at 37°C (GTH12) (Mean ± SEM: 27.2 ± 4.4), which was stabilized in the following weeks.

**Figure 3.**
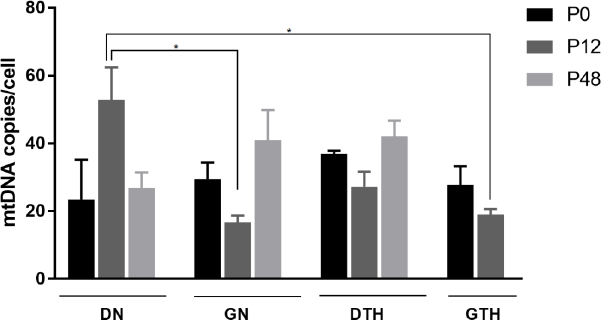
MtDNA copy number per cell. *C. albicans* SC5314 grown for 48h (P0), 12 (P12) or 48 weeks (P48) of experimental evolution. Data shown as mean ± SEM of three independent experiments in triplicate. Statistically significant differences (P < 0.05) are denoted by a and b (DN12 *versus* GN12 and GTH12). The number of mtDNA molecules/cell was determined by Real time PCR using Sybr Green with *COX2* quantification normalized by the nuclear *ACT1* gene.

**Table 2.** Mean mtDNA copy number for *C. albicans* SC5314 experimental evolution samples after 48h (P0), 12 (P12) or 48 (P48) weeks of growth. Values depicted as Mean ± SEM of three independent experiments in triplicate. Superscripts a and b denote statistically significant differences (P < 0.05) and n/a is a value not available due to the absence of yeast cells.

| Strain SC5314 | P0 | P12 | P48 |
| --- | --- | --- | --- |
| DN | 23.4 ± 11.8 | 52.7 ± 9.8<sup>a</sup> | 26.8 ± 4.6 |
| GN | 29.4 ± 4.9 | 16.7 ± 2<sup>b</sup> | 40.9 ± 8.9 |
| DTH | 36.9 ± 0.9 | 27.2 ± 4.4 | 42.1 ± 4.6 |
| GTH | 27.7 ± 5.5 | 18.9 ± 1.7<sup>b</sup> | n/a |

### Emergence of nuclear genome variants under hypoxia and heat shock

Although sequence changes were not found in mtDNA throughout *in vitro* evolution, molecular epidemiological studies show that *C. albicans* strains have high levels of intraspecific genetic diversity in its nuclear genome. Accordingly, we investigated if hypoxia and 37°C could play a role in this diversity by sequencing fragments of seven nuclear-encoded genes for the samples DN, GN, DTH and GTH (Table 1) after 12, 24 and 48 weeks of continuous growth. The gene fragments selected for sequencing were those constantly used for *C. albicans* strain typing as described in (Bougnoux et al., 2003) (MLST), namely *AAT1a, ACC1, ADP1, MPiB, SYA1, VPS13* and *ZWF1b*. Sequence variations in these MLST genes are also indicative of microvariation and adaptive processes in the nuclear DNA (Sampaio et al., 2010). Our sample SC5314 P0 showed identical MLST sequences as those deposited previously in the database (www.pubmlst.org/calbicans) for strain SC5314, with a strain type (ST) 52 and allelic profile *AAT1a* (2), *ACC1* (3), *ADP1* (5), *SYA1* (2), *VPS13* (24), *ZWF1b* (5) e *MPIB* (9). However, sequencing of this strain after *in vitro* evolution for different periods, led to the classification of samples into 3 distinct ST, with most samples remaining with ST identical to SC5314 P0 (ST 52) and two samples that showed microvariations. After 48 weeks of continuous growth, sample GVN48 exhibited loss of heterozygosity (LOH) (R > G) in position 35 of locus *ADP1*. This locus encodes a putative PDR-subfamily ABC transporter, and becomes an ST 310 with allelic profile *ADP1* (2). Sample DTH48 also showed LOH in the positions 7 (R > A), 40 (R > G), 89 (R > G) and 124 (R > G) of locus *AAT1a*, that encodes an aspartate aminotransferase, with an allelic profile *AAT1a* 8 and ST 47. As observed, both samples grown either in normoxia at 28°C or hypoxia at 37° showed nuclear but not mtDNA sequence changes through time.

Sample GTH was cultivated under the three main conditions of our interest, namely, hypoxia, 37°C and glycerol as an alternative carbon source, because dextrose availability in the human body is limited (Rodaki et al., 2009). This new strain also showed differential adaptation and yeasts were extinct from the polymicrobial culture around 24 weeks of growth (data not shown). Therefore, we decided to perform WGS of sample GTH, after 12 weeks of experimental evolution (GTH12), to identify possible variations in comparison to sample P0 (DN). Paired-end reads obtained for sample SC5314 P0 (DN) were mapped to the diploid reference sequence previously available for strain SC5314 (A22-s07-m01-r18). The fraction of the genome sequenced for sample P0 was 0.99 to 1, with an average coverage ranging from 20.85 to 40×, depending on the yeast nuclear chromosome, while the mtDNA was sequenced with an average coverage of 11,236× (Supplementary Table 1). The consensus sequence for P0 was extracted and used as a reference for sample GTH12 mapping and variant detection. Mapping of paired-end reads of sample GTH12, after removal of duplicated reads and local realignment, resulted in an average coverage of 15.9× to 26.8× for yeast nuclear chromosomes and 1,920× for its mitochondrial genome. Satisfactory fractions of chromosome sequences were obtained, ranging from 0.98 to 1.00 (Supplementary Table 1). Although the mitochondrial genome exhibited high average coverage (1,920×), and bases with coverage up to 6,299bp, we could not detect any nucleotide variation by using the probabilistic (variants with frequency ≥ 15%) or the low frequency (≤ 1%) methods, and all variants described were detected in the yeast nuclear genome. Using the probabilistic method for variant detection and strain P0 consensus sequence as a reference, we identified 979 variants in the pool of cells sequenced for sample GTH12, from which 788 were located in non-coding and 191 in coding regions. Most variants were localized in the non-coding regions (80%, 788/979) and were single nucleotide variants (SNV) (60%, 581/979), but sample also revealed multinucleotide variants (MNV) (5%, 49/9), insertions (9.4%, 92/979), deletions (22%, 215/979) and replacements (0.2%, 2/979). Gene ontology (GO) enrichment analysis revealed that alleles with variable nucleotide frequencies between sample GTH12 and P0 were involved in distinct biological processes, especially filamentous growth, pathogenesis, cellular response to starvation, drug, or biotic stimulus and cell adhesion (Figure 4). Alleles were also predicted to be involved with molecular functions such as ATP binding and ATPase activity, zinc ion, mRNA or transcription factor and enzymes binding (Figure 4). Cellular component enrichment revealed that the proteins coded by the variable alleles could be found in different cell compartments, such as the nucleus, cell surface, cytosol and mitochondrion. At least 7 genes with GO mitochondrial category showed nucleotide changes between GTH12 and P0 (Figure 4A). These nuclear encoded mitochondrial genes with sequence variability, whose proteins are predicted to act in the mitochondrial compartment, are mainly localized in mitochondrial membranes, including putative mitochondrial carriers and protein catabolic processes. We did not detect nucleotide variations in genes coding for proteins of the respiratory chain complexes.

**Figure 4.**
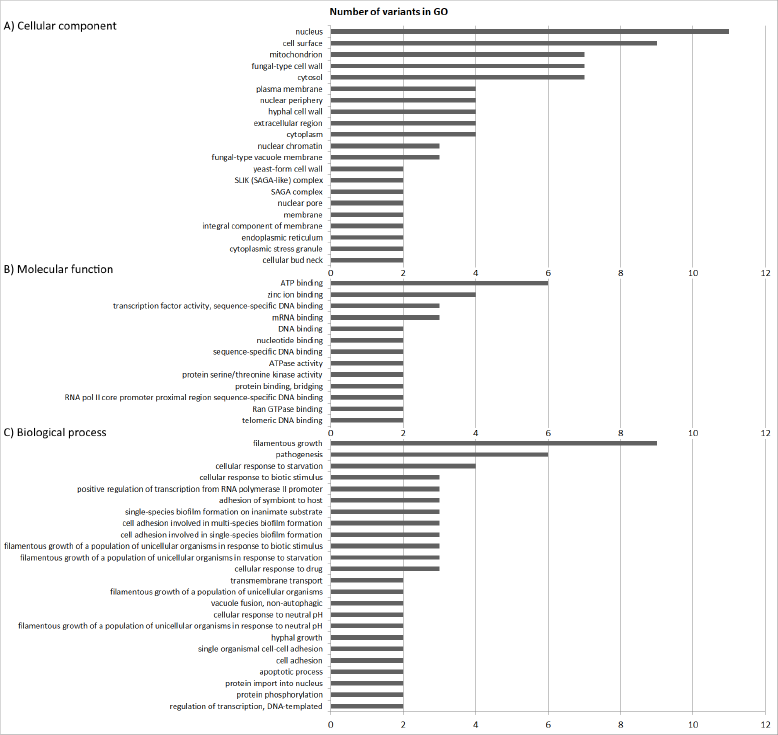
Gene ontology groups (GO) enrichment analysis of alleles with variants in sample GTH12 as compared to P0. A) Cellular component, B) Molecular function and C) Biological process.

Additionally, GTH12 variants in the coding regions were manually checked according to strain P0 genome assembly and were filtered considering their presence and read coverage in this sample, *i.e*. before *in vitro* evolution. This manual filtering resulted in at least 39 coding variants discarded due to inconsistencies and/or low-quality bases in the P0 assembly that resulted in a dubious base in the consensus sequence generated, which was subsequently used for GTH12 assembly and variant calling. After this manual checking, we observed a total of 940 variants, with 152 in coding regions. A total of 92 SNV (60.3%, 92/152), 8 MNV (5.3%, 8/152), 2 insertions (1.3%, 2/152) and 50 deletions (33.1%, 50/152) were identified in these coding regions. The synonymous variants were 31.8% (48/152), and non-synonymous variants were the majority (68.2%, 103/152) (Supplementary Table 2). This manual analysis identified 7 and 81 coding variants in the GTH12 genome that were not present in the P0 strain genome sequencing, in sites with P0 coverage > 95× or >1 and <50×, respectively. At least 64 coding variants were present in the P0 population sequenced, with P0 coverage ranging from 5 to 158× at those sites. Supplementary Table 2 summarizes the nucleotide frequency of non-synonymous variants in strain GTH12 as compared to strain P0. At least 3 alleles of GTH12 genome assembly showed variants that were not identified in P0, although those regions had high coverage (>95×) in the P0 assembly (C1_00010W_A, C3_07980C_B, CR_06380C_B). These alleles have unknown molecular function and biological processes so far. In overall, 83 alleles showed changes in the nucleotide frequencies in the strain GTH12 as compared to P0, and 71 (85%) exhibited at least one non-synonymous nucleotide variation in their sequence (Supplementary Table 2). No nucleotide changes were found in tRNA or rRNAs sequences.

### Hypoxia and heat shock affects the methylation pattern of C. *albicans* mtDNA

To date, there is no data available on *C. albicans* mtDNA methylation patterns and how host conditions may influence it. Because we did not detect any sequence changes in the mtDNA of sample GTH12, we investigated whether hypoxia and temperature could influence methylation patterns. We performed WGBS for strain SC5314 P0 and GTH12 and evaluated their mtDNAs methylome. For strain SC5314 P0, we obtained 85% of the mtDNA sequenced (34,233bp) with an average coverage of 162×, and 82% (33,151bp) of the mtDNA of strain GTH12, with 119× average coverage. Methylation levels were analyzed in three different contexts, CpG, CHH and CHG, on both strands (Figure 5A). Strain P0, which was grown under normal lab conditions, exhibited a global hypermethylation pattern, with a mean methylation/cytosine of to 99% (Figure 5B). In addition, the methylation distribution shows that approximately 97%, 96% and 94% of the mtDNA cytosines have the methylation rate equal to or greater than 95% in the CpG, CHH and CHG contexts, respectively (Figure 6A). Meanwhile, strain GTH12, showed reduced levels of methylation, with higher heterogeneity and greater variability in the methylation levels of the mtDNA with a mean methylation of 60%, including nonmethylated sites (0%) that were not present in P0 mitochondrial methylome (Figure 5B). The methylation distribution showed that only 14%, 9.1% and 12.2% of the cytosines analyzed had values equal to or greater than 90% methylation in CpG, CHH and CHG, respectively (Figure 6B and Figure 7). Fragments of mtDNA not assembled correspond to the two repetitive and inverted regions present at the two ends of *C. albicans* mtDNA, with approximately 7 kb each (Figure 8). Therefore, we observed that the growth conditions applied to the GTH12 sample influenced the methylation profile of the yeast mtDNA, indicating that the methylation in *C. albicans* mtDNA may be related to the fungus cellular response and adaptation to environmental conditions (Figure 8). Similar results were identified for another *C. albicans* strain (L757) (Supplementary Figure 1, Supplementary Figure 2). The variability in cytosine methylation percentages, with values ranging from 0 to 100% at each site, is related to the heterogeneity of the sample, which is composed of a mixture of *C. albicans* cells and / or the occurrence of several copies of the mtDNA per cell, which may have heterogeneous methylation profiles. Also, for some genes, within a predominantly methylated population, the occurrence of unmethylated copies between them is common. This indicates that methylases may have a limiting rate, which results in incomplete methylation or that the transition from active to inactive transcription occurs through a passive dilution of methylated copies during replication (Mishra et al., 2011).

**Figure 5.**
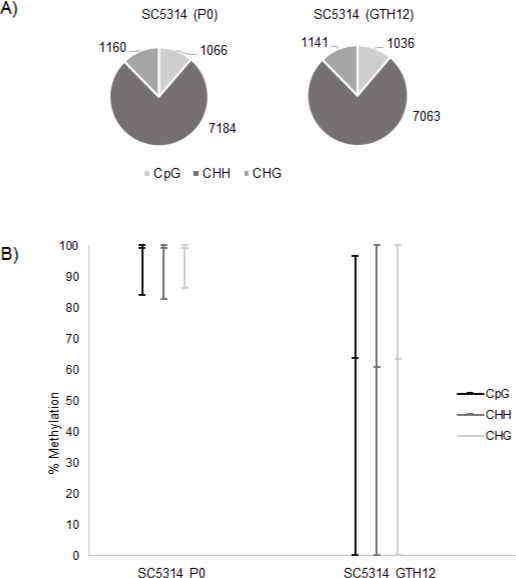
*C. albicans* SC5314 mtDNA methylation profile. A) Number of CpG, CHH and CHG analyzed on strain SC5314 mtDNA before (P0) and after (GTH12) experimental evolution. There is a prevalence of CHH sites on the yeast mtDNA, showing the importance of the methylation in non-CpG context for this pathogen. B) Percent methylation of cytosines.

**Figure 6.**
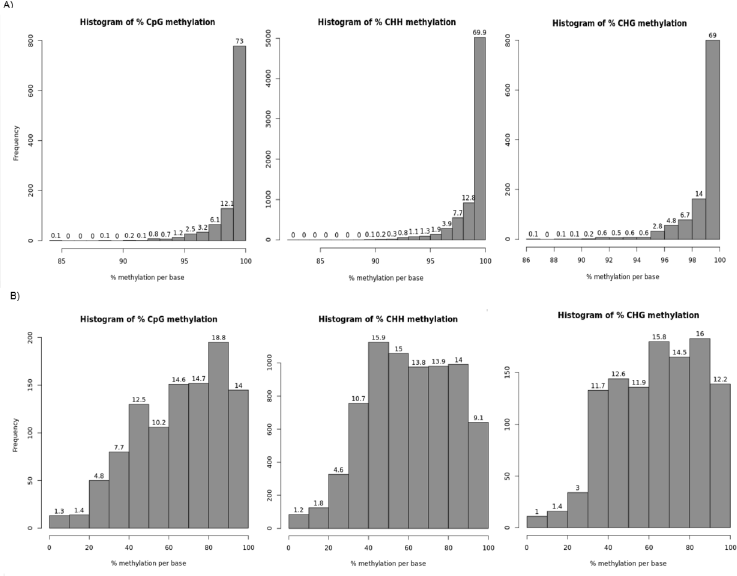
Histograms containing the distribution of the methylation percentage per cytosine for A) SC5314 P0 and B) SC5314 GTH12. The number on top of bars represent the percentage of C^met^ corresponding to the percent methylation indicated on the x-axis, while the y-axis contains the absolute frequency of C^met^. Histograms show a prevalence of hypermethylated sites for strain SC5314 P0 and a heterogeneous methylation pattern for strain SC5314 GTH12 mtDNA.

**Figure 7.**
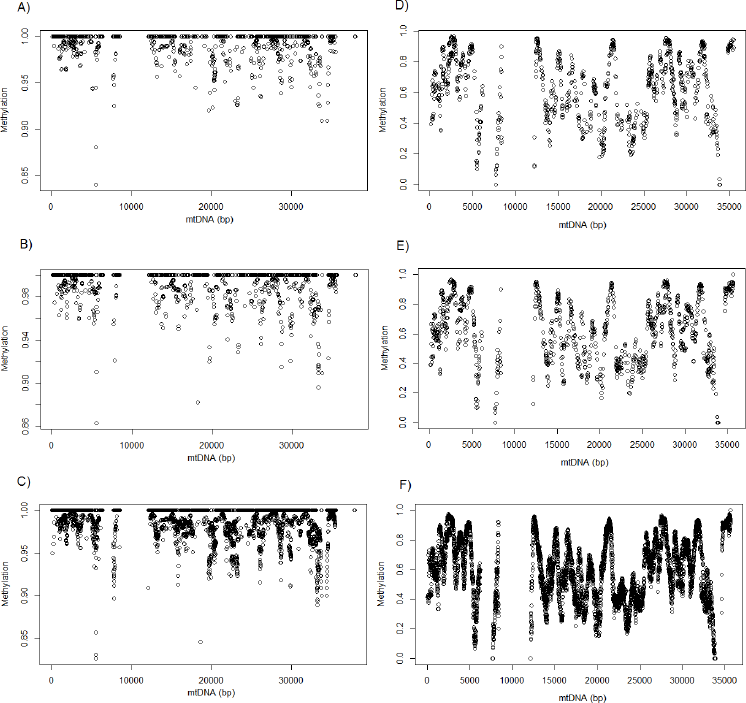
Methylation distribution throughout the mtDNA of strain SC5314 P0 (A, B, C) and GTH12 (D, E, F) on CpG (A, D), CHG (B, E) and CHH (C, F) contexts.

**Figure 8.**
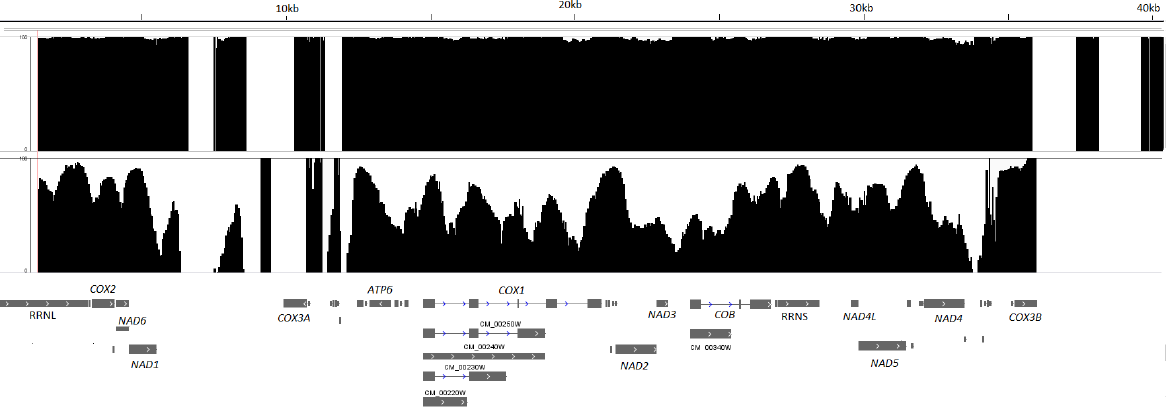
Methylation map of the mtDNA of *C. albicans* SC5314. The figure depicts the percentage of methylation per cytosine on CpG, CHH and CHG contexts for strain P0 (top row) and GTH12 (bottom row). The mtDNA fragments without a percentage of methylation correspond to the two repetitive inverted portions of *C. albicans* mitochondrial genome (5,540 – 12,382pb and 33,578 – 40,420pb).

Using the Pearson correlation coefficient, we identified that the mtDNA methylation of cytosines is not strongly correlated, with values ≤0.35 (0.35 CpG, 0.25 CHH and 0.28 CHG), suggesting environmental influence on the overall level of methylation of *C. albicans*. Global methylation profiles were also analyzed by the Spearman correlation coefficient, with similar results (data not shown).

The methylated mtDNA cytosines are mainly distributed inside introns, exons and intergenic regions (Figure 8). *C. albicans* mtDNA transcription is polycistronic, and there are only 8 promoters involved in the regulation of 8 transcription units (TU) (Kolondra et al., 2015). Among them, only TU7 promoter (CTCCTTATA), controlling the expression of *NAD4L*, *NAD5* and 3 tRNAs genes, located between positions 29,532 and 29,540, has cytosines in its sequence. For strain SC5314 P0, cytosine methylation in this promoter was close to 98%, while the values dropped to 67% for sample GTH12.

To identify differentially methylated cytosines between samples P0 and GTH12, bases with q-value <0.01 and % methylation difference >30% were considered (Akalin et al., 2012; Wang et al., 2011). When comparing SC5314 P0 and GTH12, we identified 5,520 differentially methylated cytosines, 530 in the CpG, 4,355 in the CHH and 635 in the CHG context, all of them hypomethylated in GTH12 as compared to P0. Only 1,949 cytosines (35.3%) were located in coding regions. Among genes with highest number of differentially methylated cytosines between GTH12 and P0, are RRNL, *COX1*, *COB* and *NAD4*, including possible endonucleases involved with *COX1* and *COB* splicing (Table 3). The TU7 promoter has at least 2 differently hypomethylated cytosines (29,534 and 29,535). When analyzing differentially methylated cytosines between strain SC5314 and L757, we could detect genes with cytosines hypo and/or hypermethylated between SC5314 P0 *versus* L757 P0 and SC5314 GTH12 *versus* L757 GTH12 (Supplementary Table 3), indicating that mtDNA methylation are likely strain specific and associated with the different adaptation patterns of *C. albicans* clinical isolates in the human body.

**Table 3.** Differentially methylated cytosines (C^diff met^) between samples SC5314 P0 and SC5314 GTH12. Bases with q-value < 0.01 and difference of methylation greater than 30% for cytosines with a minimum coverage ≥ 10 on both samples. All C^diff met^ were hypomethylated on sample GTH12 as compared to P0. Endonucleases coded within *COX1* (*) or *COB* (**) gene sequences.

| Differentially Methylated Cytosines ($C^{\text{diff met}}$) | | | | |
| --- | --- | --- | --- | --- |
| mtDNA localization |  |  |  |  |
| Methylation | Context | Total | Non-coding Regions | Genes (number of differentially methylated Cs) |
| hypo | CpG | 530 | 332 | RRNL (35), tRNA-Ala (5), *COX2*(2), *NAD6* (1), *NAD1* (3), CM_00230W (20)*, CM_00220W (13)*, CM_00240W (35)*, CM_00250W (51)*, *COX1* (66), *NAD2* (7), *NAD3* (2), CM_00340W (11)**, *COB* (29), RRNS (7), *NAD4L* (4), *NAD5* (6), *NAD4* (14), tRNA-Met (2), *ATP6* (6), *ATP8* (2), tRNA-Pro (3), tRNA-Gly (5) |
| hypo | CHH | 4,355 | 2,860 | RRNL (149), tRNA-Ala (7), *COX2* (36), *NAD6* (28), *NAD1* (28), *COX1* (505), CM_00220W (121)*, CM_00230W (205)*, CM_00240W (325)*, CM_00250W (437)*, *NAD2* (120), *NAD3* (32), *COB* (216), CM_00340W (131)*, RRNS (39), *NAD4L* (21), *NAD5* (79), *NAD4* (124), tRNA-Met (6), *ATP6* (66), *ATP8* (13), tRNA-Pro (8), tRNA-Cys (9), tRNA-Gly (9) |
| hypo | CHG | 635 | 379 | RRNL (47), tRNA-Ala (1), *COX2* (6), *NAD6* (4), *NAD1* (8), *COX1* (66), CM_00220W (11)*, CM_00230W (22)*, CM_00240W (39)*, CM_00250W (52)*, *NAD2* (20), *NAD3*(4), *COB* (37), CM_00340W (20)**, RRNS (6), *NAD4L* (5), *NAD5* (14), *NAD4* (28), *ATP6* (3), *ATP8* (3), tRNA-Pro (2), tRNA-Gly (2) |

## DISCUSSION

In this study we have carried out an experimental evolution scheme of *C. albicans* SC5314 where the yeast was subjected to serial passages and maintained under hypoxia and heat shock in polymicrobial cultures for up to 48 weeks. The human body is a complex environment for microorganisms, and common laboratory microbial cultivation schemes usually fail to represent important components of this system to prioritize optimal *in vitro* culture growth. To survive in the human body, *C. albicans* must find and adjust its metabolism to compatible nutrients, circumvent the host immune system and interact with other members of the human microbiome, such as *S. epidermidis* and *P. acnes*. These are other human commensals and opportunistic pathogens involved in nosocomial infections (Achermann et al., 2014; Peleg et al., 2010).

Several authors have shown high genetic variability among *C. albicans* isolates, suggesting that this opportunistic pathogen undergo adaptation and microevolution when colonizing the human body (Forche et al., 2009; Morschhäuser et al., 2000; Wartenberg et al., 2014). Although many studies investigated and characterized these intraspecific variations in the nuclear DNA, little is known about this species mtDNA, especially concerning its evolutionary rate and its effect on yeast fitness and virulence. Along with our study, several genetic changes in the nuclear DNA of *C. albicans* were already reported in short periods, after few months (1 to 4) for samples collected from patients with recurrent infections (Sampaio et al., 2010; Schröppel et al., 1994) or even after just few days (7), during a murine infection model (Forche et al., 2009). Although we did not find any sequence variation in experimental evolution samples mtDNA, their nuclear DNA exhibited microvariations that could be detected by MLST. This technique is used for strain typing and epidemiological studies concerning *C. albicans* but also as an indicative of genomic microadaptations that may be related to adaptive processes (Sampaio et al., 2010). Among the seven loci sequenced, we identified LOH in at least two of them after 48 weeks of continuous growth. Locus *ADP1*, which exhibited LOH in sample GN48, codes for a protein that is an integral component of the membrane and a putative PDR-subfamily ABC transporter, while *AAT1a*, altered in DTH48, codes for an aspartate aminotransferase which is predicted as a cellular component of the yeast mitochondrion. LOH is a common well-described phenomenon for *C. albicans*, known as one of the primary means by which this species develop intraspecific variability and population diversity (Cowen et al., 2000; Forche et al., 2005; Odds and Jacobsen, 2008). Nucleotide variants, which can be initially rare in the population, can be randomly increased by genetic drift or by selective pressure when they are beneficial to the yeast’s fitness (Ellegren and Galtier, 2016) and interfere with subpopulations frequencies in the cell sampling. Studies have shown that *C. albicans* infections can be a result of a selective proliferation of a single lineage present in the commensal population of the patient prior to an invasive infection event (Odds and Jacobsen, 2008; Odds et al., 2006). Therefore, among the sequenced population, yeasts with increased fitness and adaptation to the environmental condition tend to overgrow in comparison to other yeasts. Recurrent infections in patients may occur through the maintenance of the same isolate over time, but with the occurrence of adaptive microvariations in their genome, or the replacement of the initial strain with a more adapted/virulent one (Odds and Jacobsen, 2008; Odds et al., 2006). Mutant low-virulence strains of *C. albicans* SC5314 that are respiration-deficient were obtained during a murine *in vivo* infection model after only five serial passages. These strains, although indistinguishable from the initial SC5314 strain by molecular methods such as karyotyping and restriction enzymes, exhibited less virulence, decreased ability to kill infected mice or cause extensive damage to their internal organs, as well as lower growth rates and increased mtDNA copies and oxygen consumption through uncoupling of the respiratory chain (Cheng et al., 2007). Because sample GTH12 showed a decreased respiratory rate as compared to GTH (P0) and was grown under four important conditions also present *in vivo*, such as low oxygen, 37°C, absent dextrose and in the presence of bacteria, simulating the human microbiota, we decided to fully investigate this sample genome sequence and DNA methylation patterns, especially from its mtDNA. In addition to aerobic cellular respiration, oxygen also participates, among other things, in several cellular processes, such as iron metabolism, biosynthesis of heme groups, fatty acids, sterols and NAD (Grahl et al., 2012; Setiadi et al., 2006). Thus, it is expected that the yeast respiratory rate might influence their interaction with host niches. WGS of strain GTH12 revealed microvariations in the nuclear DNA, identified as fluctuations on the nucleotide frequencies found on strain GTH12 and P0 genome assemblies. Through our WGS experiment, probabilistic variant detection and manual filtering, we could identify several alleles with variants and nucleotide frequency altered between those samples (Supplementary table 2).

It is expected for mtDNA to have a higher mutation load when compared to nuclear DNA (around 5 to 10 times higher) (Brown et al., 1979), especially due to the reactive oxygen species production during the electron transport chain reaction, where approximately 0.2 to 2% of the oxygen is incompletely reduced to anion superoxide, making the mtDNA more prone to damage by the oxidative stress (Alexeyev et al., 2013; Balaban et al., 2005; Gao et al., 2008). Therefore, our initial hypothesis was that continuous growth of *C. albicans* under hypoxia and 37 °C, conditions found in its host, could induce adaptive changes and / or damage in its mtDNA influencing the variability of the intraspecific mitochondrial genome observed previously. The high mtDNA evolutionary rate makes this genome suitable for population and phylogenetic studies, including intraspecific analysis (Ballard and Rand, 2005; Kimura, 1983). Accordingly, our group, studying the whole mtDNA of *C. albicans* clinical isolates, identified 1.1% sequence diversity in three intergenic regions (IG1 - tRNA Gly/*COX1*, IG2 - *NAD3*/*COB*, IG3 - RRNS/*NAD4L*) (Bartelli et al., 2013). Sequencing of these intergenic regions in our experimental evolution samples during different time points (P12, P24 and P48) showed no mtDNA sequence variation as compared to sample P0. Although the Sanger method used for IGs sequencing has limited sensibility in detecting population variants under 20% frequency, the whole mtDNA sequencing, with an average coverage of 1,920.7× for sample GTH12, did not show any sequence changes in this sample grown under hypoxia and 37°C for 12 weeks. The mtDNA integrity was confirmed by Long PCR amplicons of at least two ~ 10kb regions of the 40kb mitochondrial genome (data not shown) and WGS of a second *C. albicans* strain (L757) that was subjected to the same growth conditions for the same time (L757 GTH12), also resulted in no variants detected as compared to L757 P0 mtDNA (data not shown).

Mitochondrial DNA content is variable among cells and is necessary to overcome mtDNA damage or energetic demands (Blank et al., 2008; Moyes et al., 1998). Eukaryotic cells contain multiple mitochondria that are constantly in a dynamic process of fusion-fission that enables an exchange of content and redistribution of mtDNA in face of variable environmental changes. In *S. cerevisiae*, cells contain 50 to 100 copies of mtDNA (Chen and Butow, 2005) and the variability of mitochondrial morphology, which undergoes constant fission and fusion, contributes to the distribution of mtDNA between cells (Osman et al., 2015). We identified an average of 23.4 ± 11.7 (Mean ± SEM) and 29.4 ± 4.9 mtDNA copies per *C. albicans* diploid genome, when cells were grown under normoxia for 48h (DN and GN, respectively). In *Candida glabrata*, mitochondrial fusion and fission is an important mechanism of protection against oxidative stress (Li et al., 2015). Although little is known about the mitochondrial dynamics and morphology of *C. albicans*, deficiency in mitochondrial fusion in *Fzo1* mutants triggers loss of mtDNA content and consequent decrease in its fitness, since *C. albicans* is a petite-negative yeast (Thomas et al., 2013, Calderone et al., 2015). Our results showed that *C. albicans* short (48h) and long-term exposure (12 and 48 weeks) to hypoxia and 37°C did not influence mtDNA copy number per cell as compared to strain DN (P0) (Table 2). Exposure to hypoxia and 37°C, even after 48 weeks, does not appear to significantly influence the mtDNA content (Figure 3, Table 2). Since the variation in the amount of mtDNA per cell may be related to the occurrence of damage, the fact that we did not find significant differences in the number of copies of mtDNA throughout the experimental evolution samples corroborates the data obtained for mtDNA sequencing and integrity.

Although we did not find variations in mtDNA sequences when yeasts were subjected to hypoxia and 37°C, we observed that these environmental conditions were able to affect the methylation profiles. Epigenetic factors, such as DNA methylation, can regulate the cellular response without the presence of changes in DNA sequences, and these factors can be modified and reversed by internal and external cellular stimuli, such as pollutants, oxidative stress, temperature and nutrients (Byun and Baccarelli, 2014). The induction of reactive oxygen species in human cells, for example, leads to a decrease in the mtDNA methylation, probably due to the greater compactness of the mtDNA in order to protect it, since the nucleoid protein TFAM has greater DNA binding affinity with damaged DNA (Rebelo et al., 2009). Although recent studies detected histones in the mitochondria (Choi et al., 2011), the general consensus is that mitochondrial histone complexes do not exist and that the methylation of mtDNA also plays a role in its stability and functioning (Byun and Baccarelli, 2014). In addition to being involved in energy production, mitochondria produce several epigenetic-related metabolites, such as NAD+, ATP, alpha-ketoglutarate, acetyl coenzyme A and SAM, which are necessary substrates for nuclear transcriptional and epigenetic processes, such as chromatin remodeling, histone modification and nucleosome positioning (Shaughnessy et al., 2014). In the human mitochondrial genome, the methylation pattern is uniform and constant throughout the molecule, with some differentially methylated sites between different tissues or samples collected at different time points and differential methylation at gene start sites, suggesting the existence of a regulatory mechanism that needs to be further elucidated (Ghosh et al., 2014). In this study, for the first time, we characterized the mitochondrial methylome of *C. albicans* and verified that the environmental conditions in which the pathogen is found may interfere with its methylation pattern. In our samples, methylation occurs in the CpG and non-CpG (CHH, CHG) contexts, uniformly throughout the molecule, in both gene bodies and non-coding regions. In *C. albicans*, other authors (Mishra et al., 2011) identified that the methylation of nuclear genome cytosines is also distributed throughout the gene bodies, and the presence of methylation plays a direct role in the inhibition of transcription. Meanwhile, in the human nuclear genome, several recent studies have shown that methylation in the gene body is a very frequent phenomenon and that methylation at the CpG nuclear gene promoters may not play such a large role in the gene regulation of most autosomal genes, with histone acetylation or methylation being more relevant in this context (Maunakea et al., 2010). Moreover, increased methylation along the gene sequence is related to increased transcription (Cokus et al., 2008; Flanagan and Wild, 2007; Maunakea et al., 2010; Rauch et al., 2009). In our samples, we observed a decrease in the overall methylation profile of the GTH12 sample, which may be associated with the transcriptional profile of *C. albicans* in adaptation to hypoxia, which is marked by increased expression of genes associated with glycolysis and decreased expression of genes involved in the mitochondrial tricarboxylic cycle (Krebs cycle) and oxidative phosphorylation, that are oxygen dependent (Setiadi et al., 2006; Synnott et al., 2010). The mtDNA methylation pattern and its consequences to the cell are still poorly understood and may be associated with cellular and mitochondrial responses to environmental stressors (Shaughnessy et al., 2014). Although there are few studies, mtDNA methylation is likely to influence gene expression, biogenesis and mitochondrial functions (Iacobazzi et al., 2013). In humans, the effect of mtDNMT1 (mtDNA methyltransferase 1) on mitochondrial gene transcription has been shown to be variable, depending on the strand in which the genes are encoded (H or L) and the gene in question. While overexpression of DNMT1 led to *NAD6* decreased transcription, which is encoded on the L-strand (cytosine-rich), *ATP6* and *COX1* genes, which are encoded on the H-strand (guanine-rich), did not have their transcription altered by the increase in *mtDNMT1*. However, *NAD1, which* is also coded on the H-strand, had a significant increase in its transcription. The mtDNMT1 enzyme was identified associating with mtDNA CpG sites, especially in the D-loop region, where the origin of replication and promoters are located, and also in rRNA and protein-coding gene sequences (Shock et al., 2011). It is believed that consistent with nuclear DNA, non-methylated sites in the mtDNA are caused by proteins that hinder DNMT access and consequently, cytosine methylation. In humans, DNMTs have differential access to different mtDNA sites, based on the level of proteins in their nucleoids. The ratio between the concentrations of TFAM, the major constituent protein of human nucleoids, and mtDNA may be one of the major regulators of mitochondrial activity (e.g. replication, gene expression) and may vary according to the metabolic demand of the cell. Methylation levels may be influenced by factors known to alter nucleoid structure, such as a decreased TFAM / mtDNA ratio, which leads to a less compacted mtDNA that is more accessible to DNMT, and therefore, have higher methylation rates (Rebelo et al., 2009). Therefore, unlike the common effect expected in nuclear DNA, higher levels of methylation in mtDNA may be associated with its lower compaction and, consequently, increased gene activation. In *S. cerevisiae*, Abf2 level, which is homologous to TFAM, is known to be variable in yeast cells according to its metabolic necessity. The Abf2 / mtDNA ratio is reduced under conditions that favor aerobic respiration, while their levels increase under conditions unfavorable to respiration (Xiong and Laird, 1997), suggesting an increase in mtDNA compaction under conditions unfavorable to respiration that would lead to lower methylation rates by hindering the access of DNMTs.

In this study, we observed global mtDNA hypermethylation in strain P0, which was grown under normoxia and glucose medium, and decreased methylation in yeast cells grown under hypoxia and non-fermentable medium (GTH12). In our experiments, we also observed that several sites remained with high methylation levels in GTH12 or had differential methylation between different strains (SC5314 and L757) (Supplementary Table 3). This probably occur because the distribution of DNA biding proteins, such as human TFAM, is not constant throughout mtDNA, and some regions have a more discrete decrease in methylation even with increased concentration of this protein in the cell (Rebelo et al., 2009). In *C. albicans*, nucleoid proteins, such as Gcf1 which is the most important described so far, participate in several functions, such as mtDNA replication (Visacka et al., 2009) and may have a role in the regulation of methylation of specific sites in the mtDNA to which they are associated and of the level of compactness they may induce to the molecule. When analyzing the Pearson correlation coefficients between the global methylation profiles of samples P0 and GTH12, we observed medium to low values of correlation and occurrence of differentially methylated sites between different culture conditions and strains. These results indicate that in addition to environmental conditions affecting the mtDNA methylation patterns, this response may be lineage-specific and related to adaptation and differential virulence of these strains, as previously described (Padovan et al. 2009). To our knowledge, this is the first study on *C. albicans* mtDNA methylation. The complexity of niches in the human body and microbiome species interactions could also interfere with host-pathogen epigenetic modulation. Bacteria, for example, can produce host-like molecules that are capable of post-translational modifications, such as acetylation, ubiquitination, and phosphorylation in histones and chromatin-associated proteins, in order to favor their survival (Gómez-Díaz et al., 2012). Thus, it is possible that the bacteria co-isolated and in co-culture with *C. albicans* also may influence changes in the methylation pattern of its DNA, and this phenomenon needs to be further investigated.

## MATERIALS AND METHODS

### C. albicans strain and experimental evolution

We used the *C. albicans* reference strain SC5314, kindly provided by Dr. A. Mitchell, Carnegie Mellon University, as the ancestor for the experimental evolution. Cells were grown on YPD plate (1% w/v yeast extract; 2% w/v peptone, 2% w/v dextrose, 2% w/v agar) and a single colony was used for overnight growth on YPD broth at 28°C, 150 rpm. For the experimental evolution, 1×10^5^ cells of this inoculum were displayed on tubes containing 10mL of fresh YPD or YPG (1% w/v yeast extract; 2% w/v peptone, 2% w/v glycerol) media and samples were incubated at different temperatures (28°C or 37°C) and oxygen availability conditions (normoxia or hypoxia) for 48 weeks (Table 1). Once a week, 0.1% of cell cultures (10µl) were transferred to fresh 10mL medium while a 1mL aliquot was stored in 20% glycerol at -70°C. When necessary, strain SC5314 was grown for 48 hours in normoxia at 28°C and YPD medium (P0) and experimental evolution samples (P1, P12 or P48) were grown for 7 days (from -80°C previous stock cultures) in the appropriate growth conditions. To generate a hypoxic environment, cells were maintained in hermetically closed jars with a hypoxia generator (Microaerobac, Probac do Brasil) with an atmosphere of 5-15% O_2_, while cells under normoxia were kept under 150rpm agitation with O_2_ levels close to 21% (atmospheric).

### Growth, cell viability and respiratory rate determinations

An overnight culture (28°C, 150 rpm, YPD medium) of *C. albicans* SC5314 P0 was washed twice in a sterile saline buffer and diluted to an optical density (OD_630nm_) of 0.01 in 200μL of YPD or YPG media in a 96 well microtiter plate. Plates were incubated at normoxia and 28°C or hypoxia and 37°C according to Table 1 and the OD630nm was measured at 5h, 10h and 24h intervals. Doubling times were calculated according to (http://doubling-time.com/compute.php). Measures were performed for three independent experiments in triplicate. We estimated the number of generations of the samples during the experimental evolution based on the doubling time observed for samples P1.

For yeast cell viability, experimental evolution samples were grown in their appropriate conditions (Table 1) and diluted (1:100) in a Trypan blue (Sigma) 0.2% solution (Trypan blue 0.4%: saline buffer, 1:1). Percentage of cell viability was determined by counting of viable and stained cells in a hemocytometer chamber in three independent experiments.

The respiratory rate was estimated by measuring the oxygen consumption of 2×10^6^ cells in an oxygraph (Oxygraph2k, Oroboros). In order to avoid bacterial interference in the O_2_ intake measures, yeast cultures were pre-treated with 30mg/ml ampicillin for 7 days, washed with sterile saline and incubated again for another 7 days (or 48 hours for sample P0) without ampicillin, under the appropriate conditions (Table 1). The absence of bacteria was confirmed by an attempt of 16S rRNA amplification and visual inspection of cultures under the microscope. Yeast oxygen measures were performed for 10 min under the appropriate media (YPG or YPD) and temperature (28°C or 37°C) depending on the sample analyzed. The last 5 min of readings were used for oxygen consumption rate calculations (O_2_ nmol/min) with the software Origin v7 (Origin Lab Corp). Readings were performed in triplicate for three independent experiments and statistical analysis was carried out with GraphPad Prism v7 (GraphPad Software Inc) by One-Way ANOVA followed by multiple comparisons tests (Tukey) and P<0.05 was considered statically significant.

### Yeast DNA extraction and sequencing of mitochondrial and nuclear fragments

Total yeast DNA was extracted using the protocol described previously by Wach et al, 1994. DNA integrity was checked by electrophoresis in 1% agarose gels stained with ethidium bromide when necessary. Mitochondrial fragments (intergenic regions and *COB*) were amplified using the primers described previously (Bartelli et al., 2013). The mitochondrial intergenic regions IG1 (tRNA Gly/*COX1*), IG2 (*NAD3*/*COB*) and IG3 (SSUrRNA/*NAD4L*) amplification reactions (20 μl) consisted of 10 pmol of each primer (forward and reverse), 10 μl of Phusion High-Fidelity Master Mix with HF Buffer 2× (Thermo Scientific) and 40 ng of DNA. Cycling conditions for IG1 were 98 °C for 30s followed by 30 cycles of 98 °C for 5s, 59°C for 20s, 72 °C for 15s and a final extension step at 72°C for 10 min; while for IG2 and IG3 conditions were 98 °C for 30s, 35 cycles of 98 °C for 5s, 60°C for 20s, 72 °C for 30s and extension at 72°C for 10 min. The amplification reaction for *COB* (50 μl) consisted of 10 mM dNTP, 10 pmol of each primer, 10 μl Buffer B (2 mM MgCl2), 40 ng of DNA and 2 μl Elongase Enzyme Mix (Invitrogen). Cycling conditions were 94 °C for 5 min, followed by 35 cycles of 94 °C for 40 s, 50 °C for 40 s, 68 °C for 4 min and a final extension at 68 °C for 7 min. The nuclear MLST genes were amplified according to primers described in (Bougnoux et al., 2003). Amplification reactions (20 μl) consisted of 10 pmol of each primer (forward and reverse), 10 μl of Phusion High-Fidelity Master Mix with HF Buffer 2× (Thermo Scientific) and 40 ng of DNA. For the loci *MPib, VPS13, ACC1, AAT1a, ADP1 e SYA1* cycling conditions were 98°C for 3 min, 35 cycles of 98°C 5s, (annealing temperature)°C 20s e 72°C 45s, followed by 72°C for 10 min. Annealing temperatures were 59°C for MPIb, 54°C for *VPS13*, 56°C for *ACC1* and 55°C for *AAT1a, ADP1* and *SYA1*. The locus *ZWF1b* amplification was carried out with 98°C for 3 min, 35 cycles of 98°C 30s, 50°C 20s, 72°C for 15s with an elongation step of 72°C for 10 min.

PCR amplicons were ethanol precipitated or gel purified (Illustra GFX PCR DNA and Gel Band Purification Kit, GE Healthcare) and both strands were sequenced by dideoxynucleotide chain termination method (Sanger et al., 1977) using fluorescent BigDye terminator cycle sequencing kit (version 3.1; Applied Biosystems) in an ABI Prism 3130×l automated sequencer (Applied Biosystems) according to the manufacturer’s instructions. Reads obtained were mapped to the reference sequence (sample P0) for the identification of possible nucleotide variants using Geneious v.6.1.5 (Biomatters Ltd.). Variants were filtered for considering only those with at least 3× coverage with a 25% frequency, a p-value of 10^-6^ (0.0001% probability of finding the variant by chance) and a Phred score of at least 40 (probability to be incorrect 1 in 10,000 and accuracy of 99.99%). Potential heterozygosity sites (nuclear DNA) were also inspected using the Secondary Peak Calling tool implemented in Geneious v.6.1.5 (Biomatters Ltd.), considering only overlapping peak heights with at least 90% similarity, present in both forward and reverse reads (in order to avoid strand bias and false-positives) and with a minimum coverage of 3×. Heterozygous sites were identified by the presence of two overlapping peaks in the chromatograms showing the co-amplification and incorporation of two bases in the same position that were annotated with the IUPAC (International Union of Pure and Applied Chemistry) nucleotide code. Potential variants and overlapping chromatogram peaks were also visually inspected and those present in low-quality regions were not considered.

### Sequencing of 16S rRNA and bacteria identification

Sequences of primers used for amplification of 16S rRNA genes were extracted from (Sauer et al., 2005). The amplifications consisted of 25µl containing 12.5μl of 2X My Taq Master Mix (Bioline), 0.5 uM of each forward and reverse primer and 50 ng of DNA with cycling conditions of 95°C for 1 min, 30 cycles of 95°C 15s, 58°C 15s, and 72°C 30. The DNA used for amplification were derived from yeast extraction protocol described in the previous section. Amplicons were sequenced on both strands with an ABI Prism 3130xl automated sequencer (Applied Biosystems) as described previously. Reads were *de novo* assembled with Geneious and low-quality regions in the 5’ and 3’ ends of the consensus sequence with ambiguities and low-quality base calling (Phred score <20) were removed. To identify the bacterial species, the consensus sequences were subjected to a search on the Nucleotide BLAST tool (http://blast.ncbi.nlm.nih.gov) using the 16S ribosomal RNA Database (Bacteria and Archaea) optimized for GenBank Highly similar sequences (megablast). The criteria for identifying followed the recommendations described in (Janda and Abbott, 2007), with a minimum of 99%-99.5% sequence similarity for species identification. For *Staphylococcus* spp. discrimination, the gene *tuf* was amplified with a 20µl reaction volume containing 10μl of 2X My Taq Master Mix (Bioline), 0.5 uM of each forward and reverse primer and 50 ng of DNA with cycling conditions and primers described previously (Heikens et al., 2005).

### WGS and variant calling

Total yeast DNA was extracted from samples as described previously (Wach et al, 1994), for the complete sequencing of mitochondrial and nuclear genomes by Illumina MiSeq 2×300pb method in paired-end mode. The libraries were prepared with TruSeq DNA v2 Illumina kit according to manufacturer’s technical specifications. FastQC v.0.11.4 software (Andrews, 2010) was used to evaluate sequencing quality. Trimming was performed with the CLC Genomics Workbench v.7.5.1 (Qiagen) software with a quality score limit of 0.005 and removal of 45pb and 20pb from the 5’ and 3’ ends, discarding reads lower than 25pb. Once the quality filters were approved, reads were mapped to their appropriate reference sequence and variant discovery using the same software with default parameters unless otherwise specified. Reads obtained for strain SC5314 P0 was mapped to the assembly 22 of the reference strain SC5314 (A22-s07-m01-r18) available in http://www.candidagenome.org while sample GTH12 reads were mapped to the consensus sequence obtained previously for sample SC5314 P0. Duplicated reads were removed after mapping and local realignment was carried out with the Guided Realignment tool (Force realignment to guidance-variants) implemented in CLC Genomics Workbench v.7.5.1 (Qiagen). Genome annotation was performed with Annotate with GFF file tool available on the same software using the corresponding file GFF assembly 22 SC5314 (version A22-S05-M04-r02_features_with_chromosome_sequences.gff).

Nucleotide variants in the sequenced population were detected using the Bayesian method implemented in the Probabilistic variant detection tool (frequency > 15%) with stringent quality filters to decrease the occurrence of false positives, ignoring non-specific matches and broken pairs. Reads with non-specific matches and broken pairs were ignored. Variants should have a minimum of 10× coverage and 90% probability, be present in both forward and reverse reads to avoid strand bias and require a variant count of at least 3, ignoring variants in non-specific regions and filtering homopolymeric indels.

Gene ontology enrichment analysis was performed with the file extracted from the Gene Ontology (GO) Consortium website (www.geneontology.org/page/download-annotations) version 22.02.2016 using the CLC Genomics Workbench v.7.5.1 (Qiagen) software. GO access numbers were identified in the AmiGO 2 database (amigo.geneontology.org).

### MtDNA copy number and integrity

MtDNA copy number was measured by absolute quantification using Real Time PCR with Sybr Green, according to (Amaral et al., 2007) with minor modifications. Primers were designed for amplification of 277 and 181bp fragments for the mitochondrial *COX2* and nuclear *ACT1* genes, respectively (COX2F 5′GACTCTATAGGTGAGACAATCGAA3′, COX2R 5 ′CCCACATAATTCACTACATTGACC3′), ACT1F (5′GTGACGACGCTCCAAGAG 3′) and ACT1R (5 ′CCATATCGTCCCAGTTGG 3′). *COX2* and *ACT1* amplicons were gel purified (GFX PCR lllustra DNA and Gel Band Purification Kit, GE Healthcare) and quantified in NanoVue apparatus (GE Healthcare) before serially diluted (1:10) for the standard curve construction (10-^2^ to 10^-7^ ng/uL). Total yeast DNA was extracted from samples as described previously from an inoculum containing at least 1×10^7^ cells. In order to avoid interference of bacterial DNA in the initial sample quantification, cultures were previously treated with ampicillin as described for the respiratory rate experiments. 2ng of samples or standard DNA were used in 20μL reactions containing primer forward (500nM) and reverse (500nM) and 10μL of Maxima SybrGreen 2X (Thermo Fisher). Reactions were performed in Real Time 7500 (Applied Biosystems) with an initial denaturation at 95°C for 10 min and 40 cycles of 95°C 15s, 60°C 30s and 72°C 30s. Data collection was performed during the extension step. Melting curve analysis to check specificity were performed at 95°C 15s, 60°C 1 min followed by 95°C 30 sec and 60°C 15s. Each sample was run in triplicate, including negative controls for both genes, in 3 independent experiments. In each run, standard curves were built for *ACT1* and *COX2* (10^-2^ to 10^-7^ ng/µL) and only runs with a Pearson correlation coefficient of at least 0.99 and high efficiency (90% to 110%) were considered. The number of double-stranded DNA molecules present in 1 ng of a 277bp *COX2* fragment and in an 181bp of *ACT1* was 3.34×10^9^ and 5.12×10^9^ molecules, respectively, according to the bp 650 g/mol molar mass (www.thermoscientificbio.com/webtools). MtDNA copy number per cell was calculated as the ratio *COX2* / (*ACT1* /2) molecules. Data were analyzed with GraphPad Prism v 7.6 (GraphPad Software Inc) by one-way ANOVA followed by multiple comparison tests (Tukey) between groups and P < 0.05 was considered statistically significant.

### Whole genome bisulfite sequencing (WGBS)

Total yeast DNA was extracted from samples SC5314 P0 and GTH12, using the protocol previously described (Wach et al, 1994). DNA was treated with sodium bisulfite (Zymo EZ DNA Methylation Gold kit) prior to the addition of the adapters during a preparation of the libraries, according to the Illumina TruSeq Nano kit manufacturer’s instructions. Paired-end 2×300pb runs were performed by the MiSeq Illumina method. FastQC v.0.11.4 software (Andrews, 2010) was used to evaluate sequencing quality and trimming was performed with Trim Galore 0.4.0v

(Krueger, F - http://www.bioinformatics.babraham.ac.uk/projects/trim_galore/).

Read alignment was carried out with Bismark Bisulfite Mapper 0.13.0v (Krueger and Andrews. 2011) and duplicated mapped reads removed. The percent methylation was calculated with the bismark_methylation_extractor tool implemented in the Bismark program (bedGraph files) and the R package MethylKit v. 0.9.5 (Akalin et al., 2012) for the different CpG, CHH and CHG contexts. The sites analyzed were filtered and only those with Phred score >20 and minimum 10 reads coverage were considered. Histograms with coverage distribution were checked and no duplication bias was detected. Differentially methylated cytosine calculations (Wang et al., 2011) and Pearson correlation coefficients were calculated as implemented in MethylKit v. 0.9.5 (Akalin et al., 2012). For that analysis, only sites with satisfactory quality and coverage on both samples were considered. MtDNA mappings were visualized in the Integrative Genomics Viewer v.2.3.57 (Robinson et al., 2011).

### Nucleotide sequences accession numbers

*C. albicans* sequences obtained in this study have been deposited in GenBank (https://www.ncbi.nlm.nih.gov/nucleotide/), under the accession numbers listed in the supplementary data.

## ACKNOWLEDGMENTS

The authors thank Prof. Alicia Kowaltowski (Department of Biochemistry-IQ, University of São Paulo, Brazil) for suggestions on oxygen consumption experiments and oxygraph equipment availability. This work was supported by grants to M.R.S.B from Fundação de Amparo à Pesquisa do Estado de São Paulo – FAPESP, Brazil (2013/07838-0) and Conselho Nacional de Desenvolvimento Científico e Tecnológico – CNPq, Brazil (303905/2013-1). T.F.B received a FAPESP PhD fellowship (2013/08457-0).

